# The program STRUCTURE is good at detecting contemporary hybridization

**DOI:** 10.64898/2026.11.12.766246

**Authors:** Daniel Pelletier, Ailene MacPherson, Arne O. Mooers

**Affiliations:** Biological Sciences, Simon Fraser University, 8888 University Dr, Burnaby, British Columbia V5A 1S6, Canada; Mathematics, Simon Fraser University, 8888 University Dr, Burnaby, British Columbia V5A 1S6, Canada

**Keywords:** hybridization, population structure, population genetics, admixture, gene flow, simulations

## Abstract

The program STRUCTURE is widely used in conservation genetics to detect human-caused hybridization between previously isolated populations. However, its performance at this specific task is not yet known, and methods used to interpret the model output vary widely among studies. We simulated hybridization between two populations on the timescale of real-world cases and tested STRUCTURE’s ability to correctly detect hybridization based on a simple “hybrid signal”: inferring the presence or absence of a minimum of one hybrid. Overall, this returned the correct result in 100% of scenarios when Fst between the two populations was between 0.05-0.15 when using an optimal Q-threshold of 0.10. Performance was slightly reduced when divergence was very weak (Fst = 0.01, with an optimal Q-threshold between 0.30-0.35) or when the population receiving migrants was substructured, but it still returned the correct result in 89-100% of cases, depending on simulation parameters. We conclude that STRUCTURE, paired with this hybrid signal and our recommended Q-thresholds, is an appropriate tool for detecting contemporary hybridization.

**Article summary:** Human actions may be causing previously isolated populations to begin hybridizing, which can erode valuable genetic diversity. Many researchers seek to detect recent hybridization with the software STRUCTURE, which uses genotype data to classify the ancestry of sampled individuals as either pure or hybrid. Using simulations, the authors find STRUCTURE to be highly capable of correctly detecting either the presence or absence of hybridization. They provide methodological recommendations to maximize STRUCTURE’s performance at this task, which should aid conservation geneticists working to detect cases of human-caused hybridization.

## Introduction

Anthropogenic hybridization (McFarlane and Pemberton 2019), or human-caused gene flow, is of conservation concern because it can erode the unique genetic makeup of formerly distinct populations (Todesco et al. 2016) and reduce the mean fitness of wild populations (McGinnity et al. 2003; Frankham et al. 2011). It has been investigated in the context of admixture between domesticated and wild populations of the same species (Oliveira et al. 2008; Glover et al. 2017; Ellis et al. 2018) and between pairs of fully wild populations or species (Crispo et al. 2011; Grabenstein and Taylor 2017), as a result of transport of wildlife (Perry et al. 2002), range expansion due to climate change (Garroway et al. 2010), land-use changes that increase habitat connectivity (Bleeker and Hurka 2001), or even reproductive barrier breakdowns in sympatry (Seehausen et al. 1997).

Many different methods have been used to detect contemporary gene flow. Hybrids can be identified by comparing the morphology of wild-caught individuals to putative parental populations, though these diagnostics can be unreliable (see, e.g., Sanchez-Donoso et al. 2014; Glover et al. 2017; Bartomeus et al. 2020). One can also use pre-hybridization DNA samples to establish baseline native allele frequencies to which more recent samples can be compared (Poulakakis et al. 2008). Unfortunately, such historical samples are rare, and determining the date of the earliest possible hybridization event is rarely simple. These challenges have led researchers to infer hybrid ancestry from genetic samples collected at a single timepoint using population genetic signals that are disrupted by recent mixing between populations.

A widely-used model-based inference method based on contemporary genetic samples utilizes the software STRUCTURE (Pritchard et al. 2000; Falush et al. 2003). STRUCTURE clusters individuals into a user-specified number of populations using multilocus diploid genotype data of sampled individuals. It does this by minimizing Hardy-Weinberg disequilibrium and linkage disequilibrium within each population, simultaneously estimating whether individuals are purely descended from one of those populations or represent hybrids among them. No prior knowledge of the evolutionary history of the populations or sampled individuals is needed, contributing to STRUCTURE’s popularity for studying hybridization in a wide variety of taxa (e.g., wild pigs (Gama et al. 2013), bumblebees (Bartomeus et al. 2020), Atlantic salmon (Perrier et al. 2012), honeybees (Ellis et al. 2018), wild cats (Oliveira et al. 2008), quails (Sanchez-Donoso et al. 2014), ticks (Alale et al. 2024), and oaks (Eaton et al. 2015)).

Users manually set the number of populations (K) into which STRUCTURE clusters individuals, typically estimating the “real” number of populations (selecting one value of K) through the ΔK method (Evanno et al. 2005), which aims to find the value of K (over an increasing range) at which the likelihood plateaus. From STRUCTURE’s output at the chosen K, the estimated ancestry proportions (Q-values) of each individual to each of the K populations are used to classify individuals as pure members of a population, or as hybrids between multiple populations, based on a predetermined “Q-threshold”. An early simulation study recommended Q-thresholds of 0.1-0.2 to detect pures and hybrids (Vähä and Primmer 2006), but those used in empirical studies can range between 0.05 (e.g., Bartomeus et al. 2020) and 0.25 (e.g., Suni et al. 2017).

Multiple simulation studies have shown STRUCTURE to accurately infer the ancestry of individuals across a variety of evolutionary scenarios, but its performance suffers when the populations are weakly differentiated (Vähä and Primmer 2006; Wang 2017), and when gene flow between previously separate populations has been ongoing for an extended period (Toyama et al. 2020; Ravagni et al. 2021). Additionally, one of STRUCTURE’s model assumptions is that the populations in question are internally panmictic (Pritchard et al. 2000), and simulation studies have shown that K-selection tools tend to identify only the topmost layer of hierarchical population structure (Evanno et al. 2005; Janes et al. 2017), making the detection of substructure especially challenging. Accordingly, guidelines recommend against using the software to study hierarchically structured populations (Lawson et al. 2018). However, STRUCTURE’s ability to specifically detect the presence or absence of hybridization, and its performance at that task under various scenarios of differentiation, duration of hybridization period, and population substructure has yet to be explicitly tested.

One issue in the use of STRUCTURE for detecting anthropogenic hybridization is the absence of guidelines for summarizing individual-level STRUCTURE results into a population-level binary diagnosis of hybridization/no hybridization. This is reflected in the methodological inconsistencies across studies. For example, Suni et al. (2017), using a Q-threshold of 0.25, concluded there was no evidence for hybridization between native and introduced bumblebees, despite finding individuals with Q-values between 0.25-0.75. In contrast, Bartomeus et al. (2020), using a Q-threshold of 0.05, concluded that there were hybrids in their sample, even though some presumably pure bumblebees sampled from greenhouses had Q-values between 0.05-0.95.

A simple STRUCTURE-based method to detect hybridization would be to take the classification of any individual (1 or more) from a sample as a hybrid as evidence for hybridization. We focus on this approach, hereafter referred to as the “hybrid signal”, as it matches the expectation of many users of STRUCTURE: that it correctly classifies individuals as pure or hybrid descendants of the sampled populations. Here, we test the ability of that method to correctly infer the presence or absence of hybridization through simulations of recent secondary contact between an “invader” and a “native” population. We examine how the successful detection of hybridization and the optimal Q-threshold depend on the extent of differentiation between the parental populations (Fst), the duration of secondary contact, and the presence of “hidden” population substructure.

## Results

As shown in **Figure 1** (details in **Supplementary Methods**), our simulations began with an ancestral population of 1200 diploid individuals. Following a burn-in period, the ancestral population was copied into two populations of 1200 individuals each: the “Native not-invaded” and the “Invader”. These evolved in isolation until Fst between them reached one of the four target Fst levels (0.01, 0.05, 0.1, 0.15), at which point a copy of the native population was made: the “Native invaded”. We initiated one-way migration from the Invader to the Native invaded population, and sampled SNP data from individuals at two timepoints following the start of invasion (10 and 30 generations). Each sample took 75 individuals from the Invader population and 75 individuals from either the Native not-invaded or the Native invaded population, forming contrasting “No-invasion” and “Invasion” scenario datasets with which we tested the hybrid signal’s performance. The substructured scenario simulations were run in the same way as the panmictic scenario, except that the native population was composed of 3 weakly connected subpopulations of 400 individuals each.

**Figure 1.**
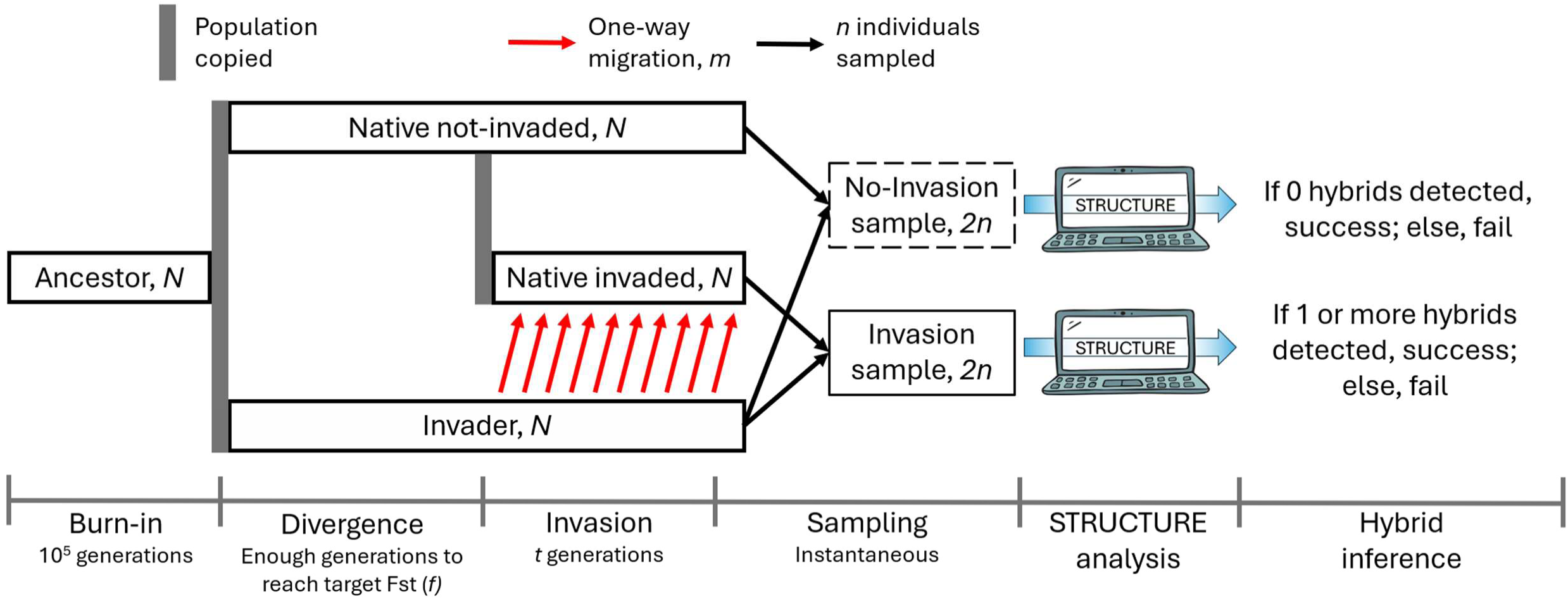
Simplified schematic of simulations and analysis. For each run, invasion starts upon reaching each of the target Fst levels (*f* = 0.01, 0.05, 0.1, 0.15), individuals are sampled after each of the invasion sampling times (*t* = 10, 30 generations), and separate “Native not-invaded” and “Native invaded” populations are simulated for the panmictic and substructured scenarios. We use 50 replicates with *N* = 1200 diploids, *m* = 10 invaders/generation, *n* = 75 individuals. For each simulation replicate where ΔK selects K=2, individuals are classified as either pures or hybrids under each Q-threshold from 0.01-0.49 (by 0.01 increments). Each STRUCTURE run is classified as a success if it correctly detects 0 hybrids in a No-invasion sample, or 1 or more hybrids in an Invasion sample; otherwise, the run is classified as a failure.

In the vast majority (797/800) of the panmictic replicates (50 replicates x 4 Fst levels x 2 sampling times), ΔK returned K=2 as the optimum number of populations (**Figure S2** and **Table S2**). The 3 replicates in which K=2 was not selected were omitted from subsequent analyses.

**Figure 2** shows how we calculated the hybrid signal’s success rate and how we identified the optimal Q-threshold ranges. At all three higher divergence levels (Fst = 0.05/0.10/0.15, at both 10 and 30 generations), success rates in the optimal Q-threshold ranges were 100% (**Figure S3** and **Table S1**), and the optimal Q-threshold ranges of those 6 scenarios overlapped at 0.10-0.14 (**Figure 3**). At Fst = 0.01, the optimal Q-threshold ranges were 0.35-0.38 and 0.30 for the 10 and 30-generation samples, respectively (**Figure 3**), with average success rates of 89% and 98% (**Figure S3** and **Table S1**).

**Figure 2.**
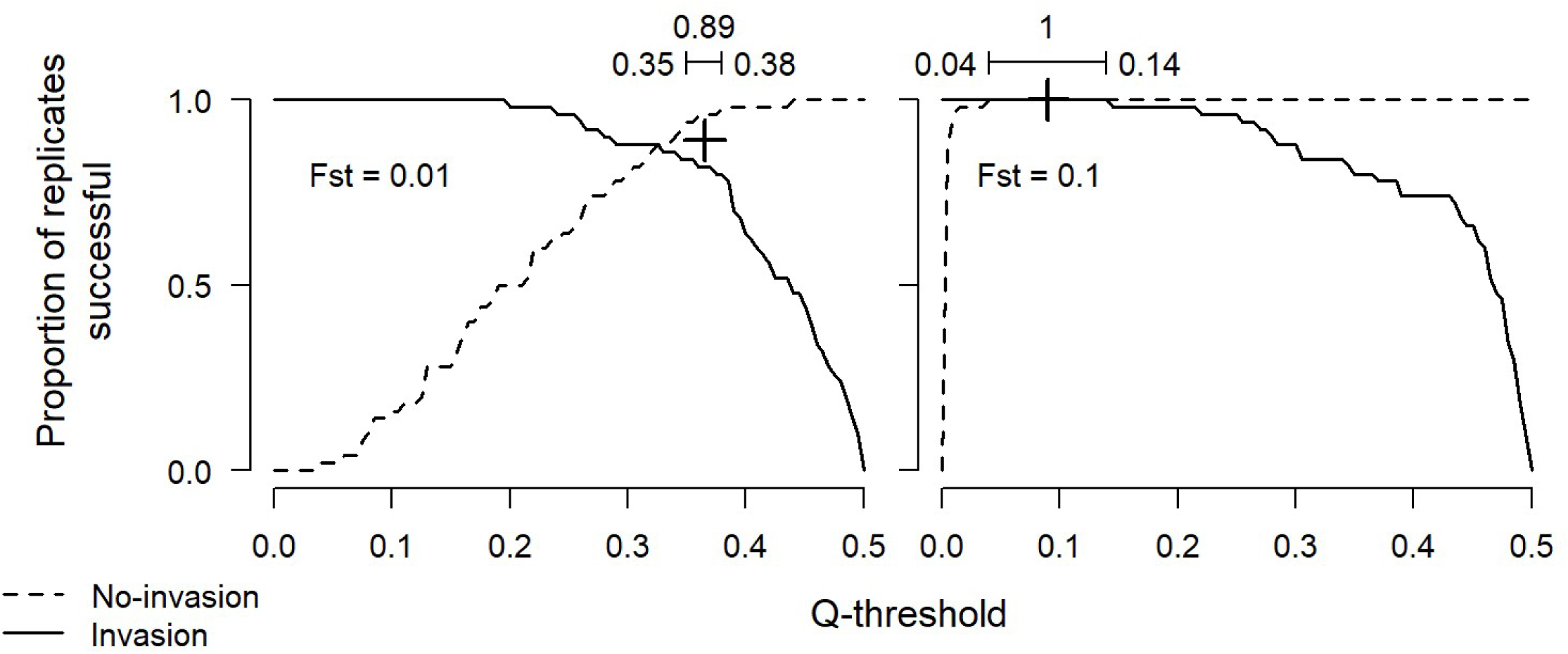
The hybrid signal’s success rate (the proportion of simulation replicates where it correctly detected hybridization/no-hybridization, averaged between the two invasion scenarios) over a range of Q-thresholds. A “No-invasion” replicate is successful if 0 individuals are labeled as hybrids; an “Invasion” replicate is successful if 1 or more individuals are labeled as hybrids. The optimal Q-threshold range (where the success rate is maximized) is indicated by the bracket at the top of each panel, with the success rate above. Results shown for panmictic scenarios at target Fst levels of 0.01 (left) and 0.1 (right), following 10 generations of invasion (all 8 panmictic scenarios shown in **Figure S3**).

**Figure 3.**
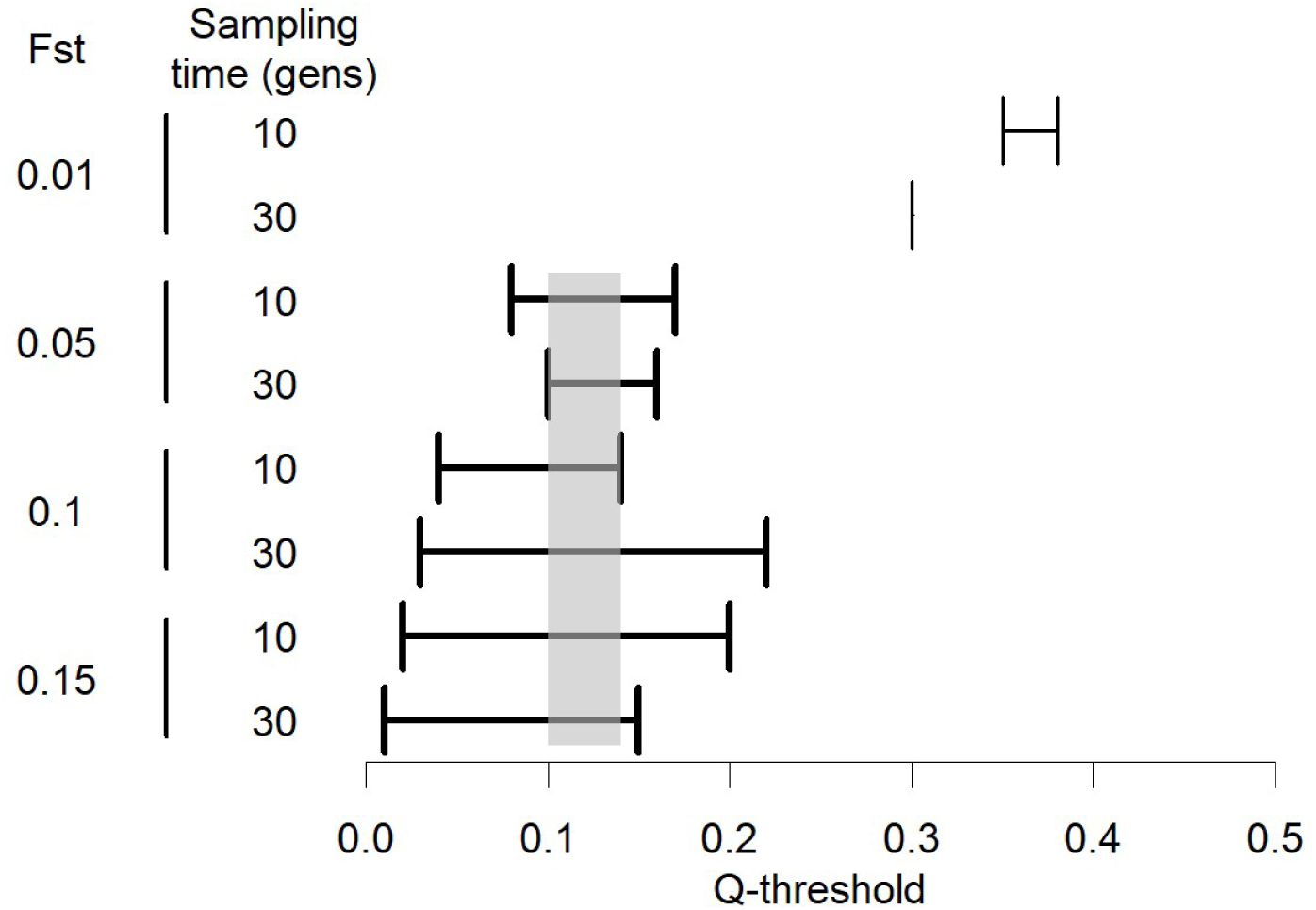
The Q-threshold ranges that maximized the hybrid signal’s success rate (averaged between the Invasion and No-invasion scenarios), for the 8 panmictic simulation scenarios. Thick bracket lines indicate scenarios in which the maximum success rate at the optimum Q-threshold was 100%. The gray shaded area indicates the range at which the optimal Q-threshold ranges of 6 scenarios overlapped.

We then calculated the “correct ID rates”: the proportion of individuals whose ancestry categories (either pure native, hybrid, or pure invader) were correctly identified by STRUCTURE, averaged across simulation replicates. When averaged between the two invasion scenarios, these tended to improve at decreasing Q-thresholds (**Figure S5**). We therefore selected the lowest Q-threshold of the optimal Q-threshold ranges identified above as our single optimal Q-threshold for each scenario: 0.10 for the scenarios at Fst = 0.05/0.10/0.15, and 0.35 and 0.30 for the Fst = 0.01 10 and 30-generation scenarios, respectively. At those selected optimal Q-thresholds, correct ID rates were 100% for all No-invasion scenarios, and ranged between 52-62% for Invasion scenarios, with higher correct ID rates at higher Fst (**Table S1**).

Finally, we evaluated the hybrid signal’s success rate when the Native population was substructured. In this scenario, ΔK selected K=2 as the optimum number of populations in only 416/800 replicates (**Figure S2** and **Table S2**). K=2 was more frequently selected at stronger divergence levels (Fst = 0.10/0.15), particularly in the No-invasion scenarios. As above, we only analysed runs where K=2 was chosen; in some low-divergence scenarios, this left as few as 2/50 simulation replicates.

Using the optimal Q-thresholds selected from the panmictic scenario, we calculated the hybrid signal’s success rate in the substructured scenario (**Figure S4** and **Table S2**). At Fst = 0.01, the success rate was slightly lower in the substructured scenario than in the panmictic one, but the low number of substructure-scenario simulation replicates for which K=2 was selected suggest these may be unreliable comparisons. Success rates at Fst = 0.05 were slightly lower than in the panmictic scenario, but still at least 90%, and success rates at the two higher Fst levels (0.10 and 0.15) were all 99-100%.

We calculated correct ID rates in the substructured scenario using the same optimal Q-thresholds (**Table S2**). In the No-invasion scenarios at the lowest Fst (0.01), the correct ID rates were 85-86% (lower than in the panmictic scenario). In the No-invasion scenarios at the three higher Fst levels (0.05, 0.10, 0.15), the correct ID rates were all 99-100%, effectively matching the panmictic scenario’s correct ID rates. In the Invasion scenarios, correct ID rates at the lower Fst levels were approximately equal between substructured and panmictic scenarios; however, at higher Fst levels, correct ID rates were higher in the substructured scenario. Additionally, correct ID rates in the substructured scenario at higher Fst levels tended to increase at the later sampling time, a trend not observed in the panmictic scenario.

## Discussion

We show that STRUCTURE, paired with our hybrid signal and optimal Q-thresholds, can reliably detect the presence or absence of contemporary hybridization between two populations. This method was successful in 100% of cases when panmictic populations were moderately to strongly diverged (Fst = 0.05-0.15), after either 10 or 30 generations of invasion, with a Q-threshold between 0.10-0.14. We recommend using a Q-threshold of 0.10 (the low end of the optimal range) because of the slight increase in the proportion of individuals whose ancestry was correctly identified. It is encouraging that we identified a single optimal Q-threshold for a wide range of scenarios, particularly because the parameters we used are typical of empirical studies’ reported Fst (e.g., Oliveira et al. 2008; Perrier et al. 2012; Gama et al. 2013; Sanchez-Donoso et al. 2014; Alale et al. 2024) and invasion timespans (e.g., Velthuis and van Doorn 2006; Glover et al. 2017).

The high success rates are likely due to the hybrid signal’s stringency rather than STRUCTURE’s infallibility, because while 100% of individuals in No-invasion scenarios were correctly categorized, only 52-62% were correctly categorized in Invasion scenarios, where most hybrids were mislabelled as pure natives (**Figure S2**). This asymmetry is reflective of our optimal Q-thresholds being conditioned on the success of a test in which not a single hybrid can be inferred in No-invasion scenarios, but only a single individual needs to be labelled as a hybrid in Invasion scenarios. Using a higher minimum number of STRUCTURE-labelled hybrids to infer the presence of hybridization could reduce this asymmetry but would require an added parameter that would depend on the true (unknown) migration rate between populations. Given the overall success of our hybrid signal, we advocate using it – i.e. a minimum threshold of 1 inferred hybrid. Q-thresholds conditioned on maximizing the correct ID rate could have avoided mislabelling so many hybrids as pures (**Figure S5**), but the optimal Q-thresholds would have been lower, resulting in the erroneous inference of hybridization in more No-invasion scenarios (**Figure S3**). This distinction is important, as previous simulation studies, including ones providing Q-threshold recommendations, have investigated STRUCTURE’s ability to correctly infer individual ancestry (Latch et al. 2006; Vähä and Primmer 2006; Sanz et al. 2009; Kalinowski 2011; Bohling et al. 2013; Wang 2017; Caniglia et al. 2020; Toyama et al. 2020; Ravagni et al. 2021), not the more specific question of whether hybridization has occurred at all.

We now briefly consider factors affecting the hybrid signal’s performance across simulated scenarios.

### Fst

As in other studies of STRUCTURE (Latch et al. 2006; Vähä and Primmer 2006; Wang 2017; Toyama et al. 2020; Ravagni et al. 2021), performance was lowest at the weakest divergence level (Fst = 0.01). At low Fst, incomplete lineage sorting and weak divergence in allele frequencies can cause STRUCTURE to mistake the genome of a hybrid individual for a pure (in the Invasion scenario) and mistake a pure individual for a hybrid (in the No-invasion scenario), leading our test to incorrectly infer the presence or absence of hybridization.

The optimal Q-thresholds were also highest in the lowest Fst scenario (0.30-0.38, vs. 0.10-0.14 at the higher Fst). This was driven by the No-invasion scenarios, in which the smaller of the two Q-values assigned to individuals were frequently quite high (**Figures S8-S9**), requiring a high Q-threshold to classify these individuals as pures. If the same Q-threshold we recommended for the higher-divergence scenarios (0.10) were used in the lowest-divergence scenario, the success rate in the Invasion scenario would be 100%, but that of the No-invasion scenario would drop to 16-48%.

### Duration of invasion

A previous simulation study of one-way invasion found STRUCTURE’s estimate of the population’s allele frequencies to deteriorate as gene flow continued (Ravagni et al. 2021). We therefore expected success rates to drop from 10 to 30 generations of invasion. However, at the three higher divergence levels (Fst = 0.05/0.10/0.15), the success rate was 100% at both 10 and 30 generations. At Fst = 0.01, the success rate even increased at the later sampling time, though this was trivially due to the No-invasion scenario, where the later sampling time simply increases Fst (**Figure S1**).

High success following 30 generations may be because STRUCTURE infers each population’s gene frequencies from the genotypes of the *sampled* individuals. In our 30-generation samples, the average native-population ancestry proportion of individuals was lower, but STRUCTURE still assigned Q-values with approximately the same distribution, labelling the individuals with the most native ancestry as pure natives (**Figures S8-S15**). This shifting baseline of what constituted the native population allowed STRUCTURE to correctly identify some hybrid individuals, despite consistently underestimating their invader-population ancestry (the same trend can be seen in Ravagni et al. (2021)). In this way, the hybrid signal correctly inferred the presence or absence of hybridization, even when STRUCTURE inaccurately inferred the gene frequencies of populations and the ancestry of individuals.

### Hidden substructure

In the substructured scenarios at low Fst, rapid but decelerating drift among subpopulations (**Figure S1**) led to ΔK most often identifying K=4 at Fst = 0.01/0.05 (**Figure S2**), though K=2 and K=3 were not uncommon. At higher divergence levels, the differentiation between the two main populations overshadowed the plateauing differentiation between the subpopulations, leading to the frequent selection of K=2. In this way, substructure can effectively be “hidden” to a user of STRUCTURE, especially at Fst in the 0.10-0.15 range. In this case, an empirical researcher would likely follow the same methodology as in a panmictic scenario, so we used the optimal Q-thresholds that were selected in the panmictic scenario. The hybrid signal was highly successful across all substructured scenarios, with near 100% success for the two highest levels of divergence, where hidden substructure was most common. Substructure may weakly hinder performance at Fst = 0.01, but sample size and confidence are low.

The most notable difference between the panmictic and substructured scenarios was at the Fst = 0.05 divergence scenario, where the success rate was reduced from 100% in the panmictic scenario to 96% (10 generations) and 90% (30 generations) in the substructured scenarios. Given the moderate frequency of K=2 results at this Fst (54/200 total replicates), this is a level of divergence at which substructure may commonly be hidden, but still somewhat negatively affects STRUCTURE’s performance at detecting hybridization. While previous studies have alluded to problems with substructure in STRUCTURE (Evanno et al. 2005; Jombart et al. 2010; Janes et al. 2017), we believe this is the first simulation study to identify this particular issue.

The mechanism may be that when the populations are weakly diverged (relative to divergence between subpopulations), running STRUCTURE with K=2 causes some of the genetic variation between subpopulations to be attributed to the Invader population, as a result of its larger sample size (Kalinowski 2011). Indeed, at Fst = 0.05, STRUCTURE overestimated many individuals’ Invader ancestry proportions, even labelling many pure natives as hybrids (**Figures S10-S11**), causing the hybrid signal to fail in some substructured No-invasion scenarios.

Surprisingly, in the Invasion scenarios at the higher Fst levels (0.10/0.15), the correct ID rates tended to be higher in the substructured scenario than in the panmictic scenario (**Table S2**). In both cases, all mistakes were true hybrids mislabelled as pure natives. It is unclear whether the improved correct ID rate in the substructured scenario was due to STRUCTURE correctly detecting the hybrid proportion of more individual’s genomes, or if, as in the No-invasion case, it incorrectly attributed some of the substructure-derived variation to invader-population ancestry, leading it to correctly detect more hybrids.

## Data availability

All scripts used for simulations, running STRUCTURE, and results analysis, as well as the main results file (including seeds to replicate simulations and STRUCTURE runs), are currently in this GitHub repository: https://github.com/danielpelletier116-prog/hybrid_detection_STRUCTURE

This will be archived in a public repository (Dryad) upon paper acceptance.

## Acknowledgements

We are particularly grateful to Darren Irwin for his close reading of an early version, saving us from ourselves. We thank members of the Hart, M’Gonigle and Mooers labs at SFU for feedback, and Sabrina Taylor for introducing AOM to STRUCTURE.

## Study funding

This study was funded by NSERC Canada (through Discovery Grants to AOM and AM, and a CGS-M 2022 award to DP), Simon Fraser University internal and donor-funded awards (Sharon Clements Biological Sciences award), and the Maud Menten Institute (2025/26 Research Accelerator Award).

